# A High-Resolution Method for the Systematic Detection of EMS-Induced Mutations in a Sequenced Population

**DOI:** 10.1101/2021.10.13.464286

**Authors:** Jared M. Simons, Tim C. Herbert, Coleby Kauffman, Marc Y. Batete, Andrew T. Simpson, Yuka Katsuki, Dong Le, Danielle Amundson, Elizabeth M. Buescher, Clifford Weil, Mitch Tuinstra, Charles Addo-Quaye

## Abstract

The precise detection of causal DNA mutations is very crucial for forward genetic studies. Several sources of errors contribute to false-positive detections by current variant-calling algorithms, and these impact associating phenotypes with genotypes. To improve the accuracy of mutation detection we propose and implemented a high-resolution binning method for the accurate detection of likely EMS-induced mutations in a sequenced mutant population. The approach also incorporates a novel clustering algorithm for detecting likely false negatives with high accuracy. *Sorghum bicolor* is a very valuable crop species with tremendous potential for uncovering novel gene functions associated with highly desirable agronomical traits. We demonstrate the precision of the proposed method in the detection of likely EMS-induced mutations in the publicly available low-cost sequencing of the M_3_ generation from 600 sorghum BTx623 mutants. The method detected 3,274,606 single nucleotide polymorphisms (SNPs) of which 96% (3,141,908) were G/C to A/T DNA substitutions, as expected by EMS-mutagenesis action. We demonstrated the general applicability of the method, and showed a high concordance, 94% (3,074,759) SNPs overlap between SAMtools-based and GATK-based variant-calling algorithms. We also implemented a novel clustering algorithm which uncovered evidence for an additional 223,048 likely false-negative shared EMS-induced mutations. The final 3,497,654 SNPs represents an 87% increase in SNPs detected in the previous analysis of the sorghum mutant population. Annotation of the final SNPs revealed 10,263 high impact and 136,639 moderate impact SNPs, including 7,217 stop-gained mutations, and an average of 12 stop-gained mutations per mutant. We have implemented a public search database for this new genetic resource of 30,285 distinct sorghum genes containing medium or high impact EMS-induced mutations. Seedstock for a select 486 of the 600 described mutants are publicly available in the Germplasm Resources Information Network (GRIN) database.

## Introduction

Plant forward and reverse genetics studies play a vital role in harnessing the untapped potential of plants in the search for new sources of fuel energy, improving crop productivity, discovering desirable agronomic traits, and enhancing ecosystem services (1-11). Despite the tremendous progress in plant gene function studies, a significant number of genes found in plant genomes still lack experimentally determined function. The rapid advances in cost-effective, high throughput sequencing techniques such as next-generation sequencing (NGS) technologies (12,13), and modern mutation breeding techniques, such as TILLING (4), had been pivotal in recent large-scale mutant studies that have provided expansive resources for crop forward genetics (14-26). A diverse collection of physical mutagens (including fast neutrons, carbon ion beam, X-rays, gamma rays and ultraviolet light) and chemical mutagens (including ethyl methanesulfonate [EMS], (N-ethyl-N-nitrosourea [ENU], and N-methyl-N-nitrosourea [MNU]) are available for inducing mutations in living organisms (27-33). A disadvantage of traditional mutagens is the modification of random genomic positions, which results in the uncertainty of genomic consequences. Although precise genome-editing techniques are on the ascendency, their general utility is still limited (34-37), and hence traditional mutagenesis techniques still remain useful. Various variant-calling algorithms are currently available for identifying induced mutations from next-generation sequencing datasets (38-42). The detection accuracy of these *in silico* tools are impacted by the inherently higher sequencing error rate of NGS technologies and other sources of error such as reference genome mis-assembly, imprecise sequence alignments and sample and library preparation errors (43). To expose false positives, present in the initial variant-calling, various practices such as the removal of shared variants, low phred-scale variant-quality score variants, extreme strand-biased variants and excessively high-coverage depth variants had been proposed (43-48). Although these filtering approaches are very potent, a possible unintended consequence is the elimination of bona fide false-negative variants. In the era of rapid advances in high-throughput phenotyping (phenomics) for both in-field and greenhouse experiments (3, 49-55), precisely detecting true variants, while filtering false-positive variants and recovering bona fide false-negative variants would greatly improve the precision of mutation detection, and greatly enhance phenotype to genotype association (52).

*Sorghum bicolor* is a grass species which is very closely related to maize and has a relatively compact genome size of around 730 million bases (56-59). Sorghum is globally cultivated for food (grain sorghum), feed (forage sorghum), beverage and fuel (cellulosic sorghum), and is characterized by the ability to grow in nutrient-poor soils (60-67). In this article we describe and implement a high-resolution approach to detect likely EMS-induced mutations in low-coverage NGS sequencing of mutant populations. This generally applicable method leverages the well-described action of EMS mutagenesis, which results in predominantly G/C to A/T DNA transitions (68,69). We also implemented a clustering algorithm for uncovering bona fide false-negative shared variants in sequenced mutant populations. Applying the method, we describe a highly improved indexed mutant collection and genetic resources for the publicly available M_3_ generation from 600 EMS-mutagenized sorghum individuals.

## Results

### NGS Mapping and Initial Variant-Calling

We implemented a high-resolution variant-calling and filtering pipeline to detect likely EMS-induced mutations in a sequenced mutant population (Figure 1). A total of 30,027,070,132 NGS paired-end reads were retrieved from the NCBI short reads archive (SRA) with an average of 50,045,117 reads per sample. Of these 98.3% (29,521,006,672) mapped to the current sorghum reference genome (version 3.1) and 28,132,018,400 were properly paired mapped reads (Table 1). Initial variant-calling using SAMtools detected 85,940,118 variants including 82,645,413 single nucleotide polymorphisms (SNPs). The percentage of SNPs that were G/C to A/T substitutions was 36.4% (30,096,471 SNPS; Table 2). Similarly using GATK variant-calling algorithm with single individual genotyping and GATK with joint genotyping, 89,338,788 and 106,488,567 variants were initially detected respectively, and these included 84,193,419 and 95,488,475 SNPs, respectively. The corresponding percentages of GATK SNPs that were G/C to A/T substitutions were 37.3% (single individual genotyping) and 37.9% (joint genotyping) respectively (Table 2). From here we simply refer to GATK variant-calling for single mutant individual genotyping as GATK, and GATK analysis with joint genotyping for all 600 mutant individuals as GATK-Joint.

**Figure 1.**
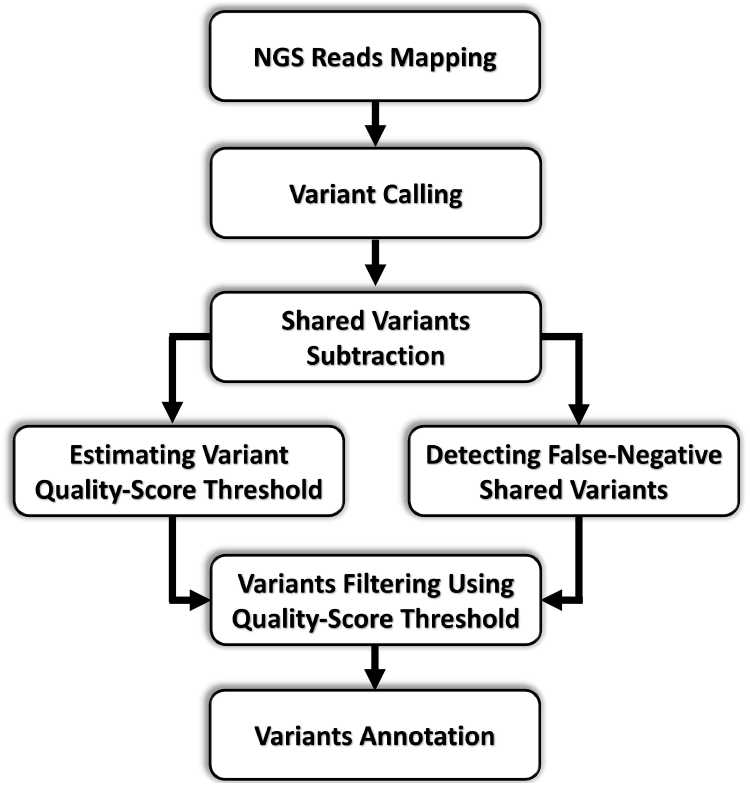
The proposed high-resolution computational pipeline for the detection of likely EMS-induced variants in the mutant population.

**Table 1.** Illumina NGS Paired-end reads mapping statistics for the 600 EMS-mutagenized sorghum BTX623 individuals.

|  | **Total** | **Mean** |
| --- | --- | --- |
| **All NGS Reads** | 30,027,070,132 | 50,045,117 |
| **Mapped Reads** | 29,521,006,672 | 49,201,678 |
| **Properly Paired Reads** | 28,132,018,400 | 46,886,697 |
| **Number of Mutants** | 600 | - |

**Table 2.** Summary variant-filtering statistics for the 600 EMS-mutagenized sorghum BTX623 individuals.

|  | **SAMtools (Sb_v2.1)<sup>a</sup>** |  | **SAMtools** |  | **GATK-Single<sup>b</sup>** |  | **GATK-Joint<sup>c</sup>** |  |
| --- | --- | --- | --- | --- | --- | --- | --- | --- |
|  | **Total** | **GCAT(%)<sup>d</sup>** | **Total** | **GCAT(%)** | **Total** | **GCAT(%)** | **Total** | **GCAT(%)** |
| **Initial Variants** | 65,384,440 | 33.7 | 85,940,118 | 36.4 | 89,338,788 | 37.3 | 106,488,567 | 37.9 |
| **Initial SNPs** | 60,407,348 | 33.7 | 82,645,413 | 36.4 | 84,193,419 | 37.3 | 95,488,475 | 37.9 |
| **Unique SNPs<sup>e</sup>** | 6,313,576 | 65.9 | 6,373,258 | 65.4 | 3,886,456 | 87.8 | 3,750,600 | 89.5 |
| **Filtered SNPs<sup>f</sup>** | 3,289,616 | 96.7 | 3,274,606 | 95.9 | 3,514,942 | 92.4 | 3,435,789 | 93.5 |

<sup>a</sup>SAMtools SNPs-calling using Sorghum BTx623 reference genome version 2.1.
<sup>b</sup>GATK SNPs-calling with single individual genotyping.
<sup>c</sup>GATK SNPs-calling with joint genotyping.
<sup>d</sup>Percentage of SNPs which are G/C to A/T DNA substitutions.
<sup>e</sup>Subset of SNPs detected only once in the entire mutant population.
<sup>f</sup>SNPs retained after SNP quality-score filtering.

### SNPs Filtering

For independently mutagenized individuals it is highly unlikely for an EMS-induced mutation to be detected in two or more mutants. The subtraction of shared (likely false-positive) variants retained 6,373,258, 3,886,456, 3,750,600 SNPs for SAMtools, GATK and GATK-Joint, respectively. Interestingly for both GATK-based approaches 88 and 90% (3,412,446 and 3,356,788) of the retained unique (non-shared) SNPs were G/C to A/T substitutions (Table 2), while the percentage for SAMtools was only 65.4% (4,166,924 SNPs). The ranges of phred-scaled SNP quality-scores for the mutations detected were 3.0 -228.0 (for SAMtools), 10.0 - 214,334.8 (for GATK) and 10.0 - 65,576,475.2 (for GATK-Joint). Visual inspection of the mutation spectrums for all variant-quality score bin categories (Figure 2, Supplemental Figures S1 and S2) shows that in all three approaches, the percentage of SNPs which were G/C to A/T transitions increased with increasing SNP quality-score values. As shown in Figure 2, for SAMtools, the G/C to A/T percentage showed a striking increase from 40.91% (at quality-score=11) to 63.73% (at quality-score=12). Similarly, for GATK and GATK-Joint the G/C to A/T percentages jumped to 64.04% (at quality-score=26) and 66.36% (at quality-score=28) respectively (Supplemental Figures S1 and S2). Based on these observations, the following SNP-quality score thresholds were selected for subsequent quality-score based filtering: 12 (for SAMtools), 26 (for GATK) and 28 (for GATK-Joint). Applying the selected SNP quality-score thresholds, 96% (3,141,908 out of 3,274,606) of SAMtools SNPs, 92.4% (3,248,054 out of 3,514,943) of GATK SNPs and 93.5% (3,211,794 out of 3,435,790) of GATK-Joint SNPs retained were G/C to A/T transitions (Figure 3; Table 2) and these are most likely EMS-induced mutations. Comparison of the detected likely EMS-induced SNPs showed a consensus (intersection) of 3,074,759 SNPs detected by all three approaches (Figure 4). The high concordance of SNPs between the SAMtools- and GATK-based algorithms suggests the filtering method is generally applicable. A total of 181,432 SNPs were SAMtools-specific, while 77,957 and 14,981 SNPs were respectively GATK- and GATK-Joint-specific. A total of 344,930 SNPs were detected in both GATK-based approaches but not by SAMtools (Figure 4). Examination of the GATK-based SNPs that were not detected by SAMtools, 55% (232,234 out of 422,887) of GATK SNPs and 58% (207,491 out of 359,911 SNPs) of GATK-Joint SNPs were G/C to A/T substitutions (Supplemental Table S1, Supplemental Files S1 and S2). Hence it is presumable that the vast majority of these GATK-specific SNPs may not be true EMS-induced SNPs. In our previous analysis of the 600 mutants, where the SAMtools SNP calling was based on version 2.1 of the sorghum reference genome, we detected 1,753,403 SNPs of which 96.7% (1,695,973 SNPs) were G/C to A/T substitutions (16). Hence the 3,274,606 SNPs detected using the currently proposed method represents an 87% (1,521,203 SNPs) increase in the number of SNPs predicted. It is possible the increase in SNPs detected could be due to differences in reference genome versions used in the separate studies. As a control, we repeated the analysis using the current approach with the older reference genome version 2.1. The initial number of SAMtools SNPs predicted in the genome version 2.1 was 60,407,348 and these included 6,313,576 unique SNPs (Table 2). Similarly, at a SNP-quality score threshold of 12, 96.7% (3,182,253 out of 3,289,616 SNPs) were G/C to A/T substitutions (Figure 3; Table 2). Hence, the increase in the number of SNPs predicted is the result of a superior method and not due to an improvement in genome assembly. Finally, we investigated the mutation spectrum for SNPs associated with extreme strand-biased reads mapping at the corresponding SNP position in the genome. Of the 307,434 strand-biased SNPs where nearly all the mapped reads (95%) at the SNP position were derived entirely from only the forward or only the reverse strand, over 94% (290,110 SNPs) were G/C to A/T substitutions. (Figure 5; Supplemental Table S2; Supplemental File S3). These supposedly extreme strand-biased SNPs are most likely EMS-induced mutations lacking sufficient read depth, because of the low-coverage (6X) sequencing of the mutant genomes.

**Figure 2.**
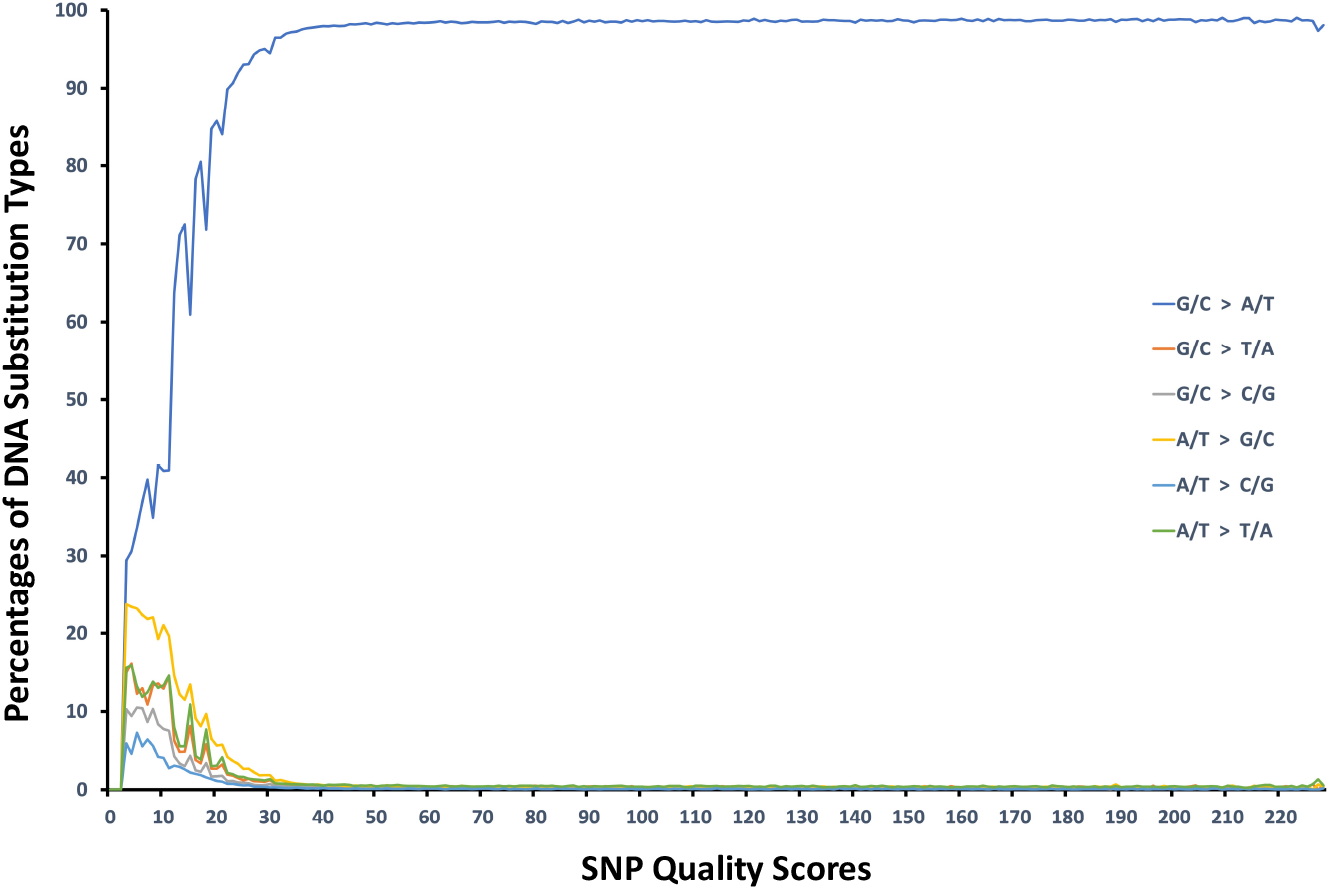
The binning approach to computing the variant quality-score filtering threshold for the initial SAMtools-predicted mutations in the 600 sorghum BTx623 mutants. The initially predicted SNPs are assigned to bin categories corresponding to their variant-quality score. The EMS mutation spectrum for all SNPs in each bin category is calculated. At a variant-quality threshold of 12, the percentages of non-G/C to A/T DNA substitutions decrease drastically, while G/C to A/T mutations become the dominant mutation type.

**Figure 3.**
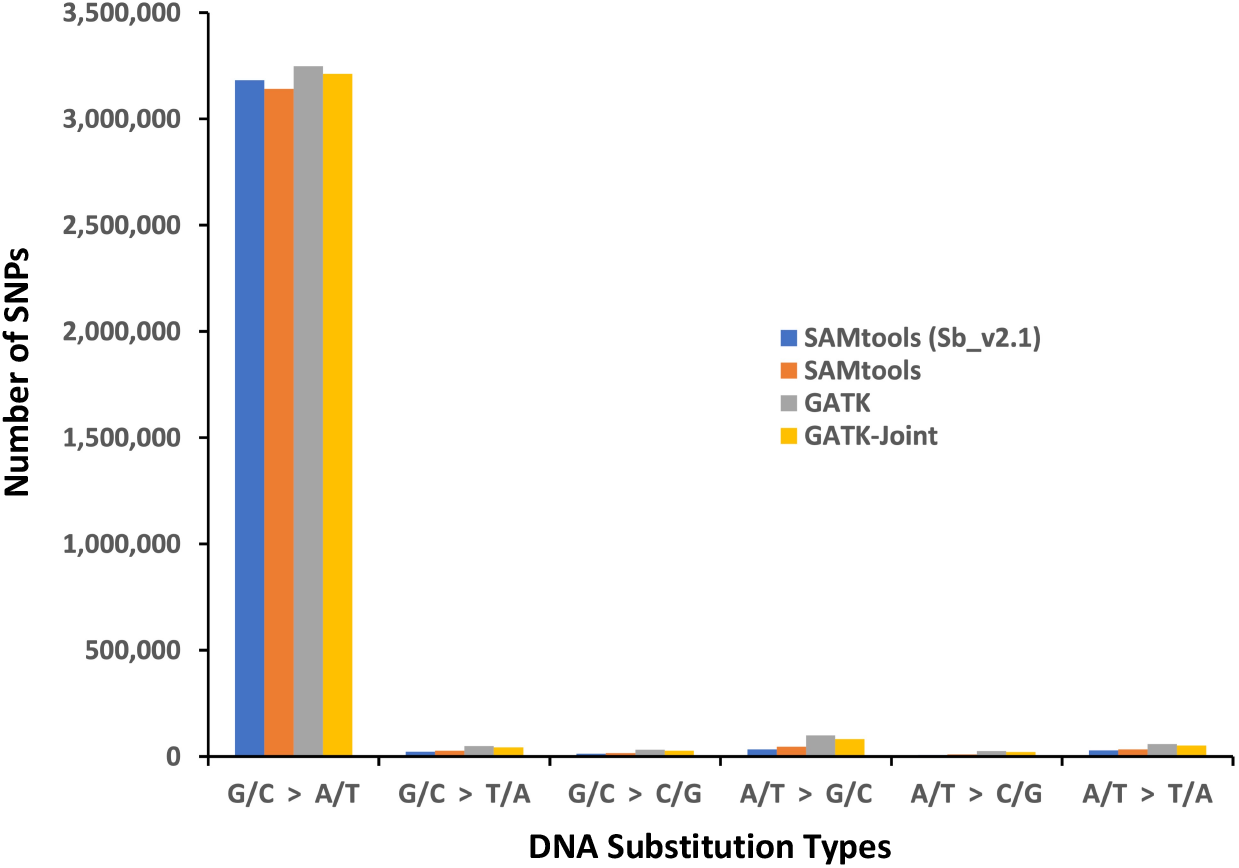
The mutation spectrums for the likely EMS-induced SNPs detected in the 600 sorghum BTx623 mutants. Using the SAMtools variant-caller and sorghum reference genome versions 2.1 (shown in above figure as Sb_v2.1) and 3.1, 96.7% and 96% of detected SNPs were G/C to A/T mutations, respectively. Using sorghum reference genome version 3.1 and GATK with single individual genotyping and joint-genotyping 92.4% and 93.5% of SNPs were G/C to A/T mutations, respectively.

**Figure 4.**
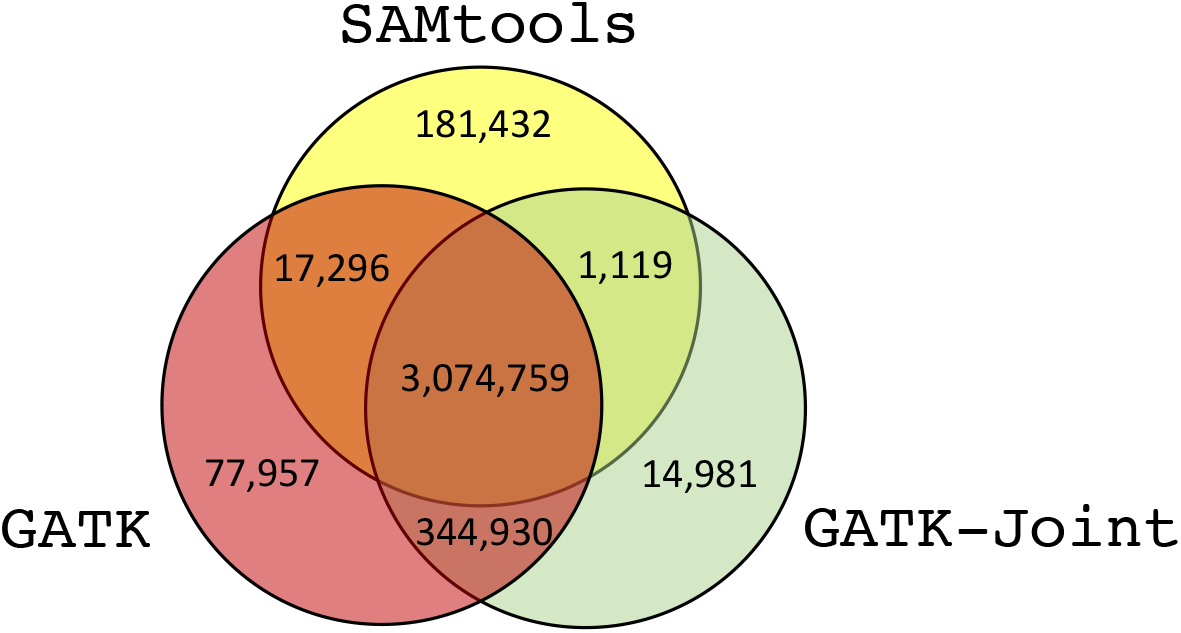
The high concordance of high-quality SNPs detected by the different variant-calling approaches, in the 600 sorghum mutants. A total of 3,274,606, 3,514,943 and 3,435,790 filtered SNPs were predicted by SAMtools, GATK (using single individual genotyping), and GATK (using joint genotyping; GATK-Joint) respectively, and 3,074,759 of these SNPs were found by all three approaches.

**Figure 5.**
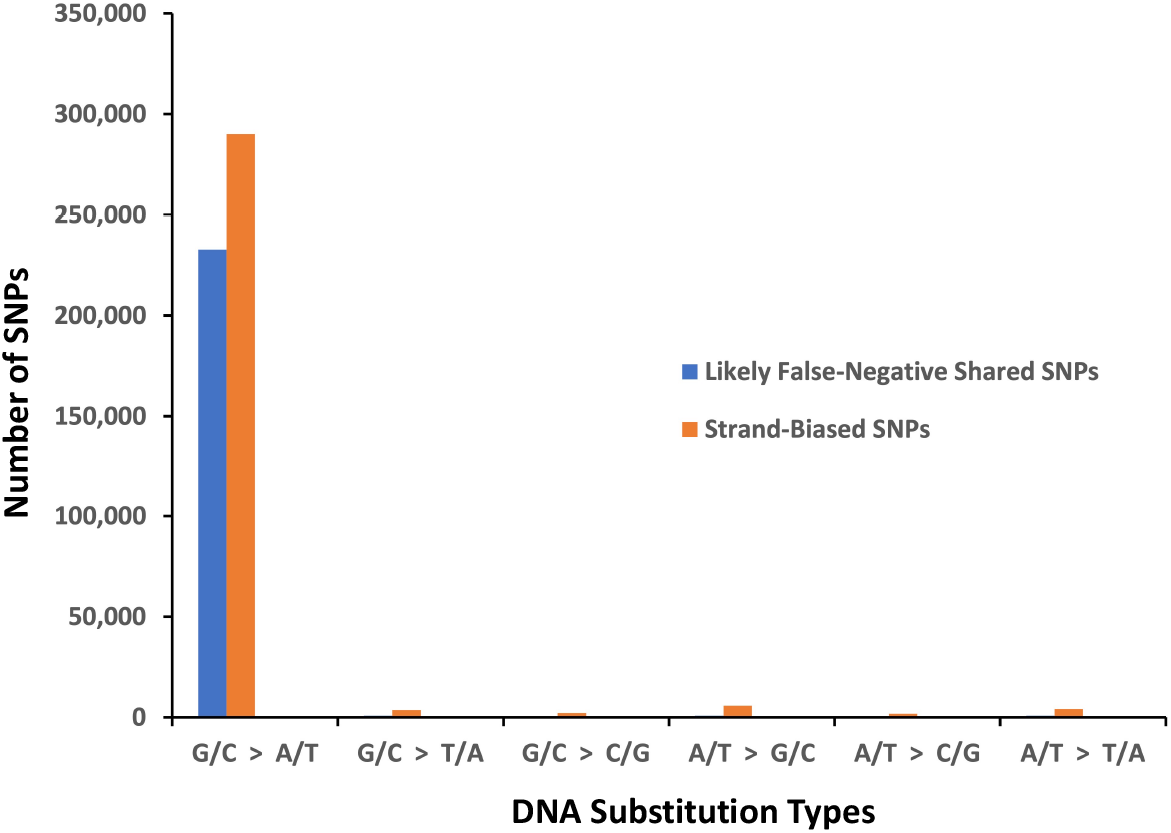
The mutation spectrums for the SAMtools-predicted extreme strand-biased SNPs and the likely false-negative shared SNPs detected by the clustering algorithm. Almost 99% (232,662) of the 235,736 likely false-shared SNPs and 94% (290,110 SNPs) of the 307,434 extreme strand-biased SNPs were G/C to A/T substitutions, respectively.

### Discovery of Likely False-Negative Shared Variants

For multiple independently mutagenized individuals in a population, the pragmatic and very effective technique of subtracting shared variants could result in the unintended consequence of the removal of false-negative shared variants. The false-negative shared variants could originate from pollen contamination, sample mislabeling, and other sources of sample or library preparation errors. Using the initial unfiltered SAMtools-predicted SNPs, we categorized all SNPs based on the number of mutant individuals they were detected in, and then computed the mutation spectrum for each category. A comparison of the G/C to A/T transition percentages for the various SNPs categorizes indicated a non-trivial signal for likely EMS-induced SNPs shared by two or three mutants (Supplemental Figure S3). We next analyzed the initially subtracted shared SNPs from the earlier SNPs filtering step, for evidence of true EMS-induced mutations. Using a clustering algorithm we implemented, we clearly distinguished between localized clusters of mutant individuals with shared mutations (likely false-negatives) and very large clusters of mutant individuals with global or near-global shared variants (likely false-positives). Overall, we identified 930,352 non-empty clusters with an average 78 mutant individuals per cluster, and an average of 1.27 shared mutations per cluster (Supplemental File S4). Ranking the clusters by the number of shared mutations within a cluster, the top 43 clusters (Figure 6) contained an average of 2,831 shared mutations, and the highest ranked cluster (*clust-39453_2*) contained 6,837 mutations shared by two mutant individuals. Most of the top-ranking clusters (41 out of 43) were of member size two (i.e., contained mutations shared by only two mutant individuals). Of the remaining two clusters, one cluster (*clust-930352_600*) contained 4,487 mutations found (or globally shared) in all 600 mutants, while the final of the top 43 clusters contained 1,238 mutations shared by a specific group of three mutants (*clust-125781_3*). Excluding the mutations from the global cluster *clust-930352_600*, the mutation spectrum for each set of recovered shared mutations for the 77 impacted mutants showed G/C to A/T percentages ranging from 96.7%-99.7% (with an average of 98.7%), and that suggests these are likely false-negative (shared) EMS-induced mutations (Figure 5). On the contrary, only 17.61% (473,979 of 2,692,200) of the SNPs originating from the 4,487 genomic positions detected in all 600 mutants were G/C to A/T substitutions. In total, we recovered a total of 235,736 likely false-negative shared SNPs originating from 117,249 distinct genomic positions, and these include 223,048 SNPs (∼95%) having a SNP quality-score of at least 12 (Supplemental File S5). This shows that, of the initially subtracted shared SNPs, 35% (232,022 of 661,156) of shared SNPs found in pairs of mutants, and only 1% (3,714 of 291,966) shared SNPs initially predicted in triplet of mutants, are likely bona fide EMS-induced mutations. Including the recovered, likely false-negative shared SNPs with variant quality-score of at least 12, we retained a total of 3,497,654 likely EMS-induced SNPs for further downstream analysis. The final retained SNPs consisted of 2,157,409 homozygous and 1,340,245 heterozygous mutations. Overall, the final SAMtools-predicted SNPs represent an average of 5,829 SNPs per mutant, and about one mutation per 125kb in the sorghum BTx623 genome.

**Figure 6.**
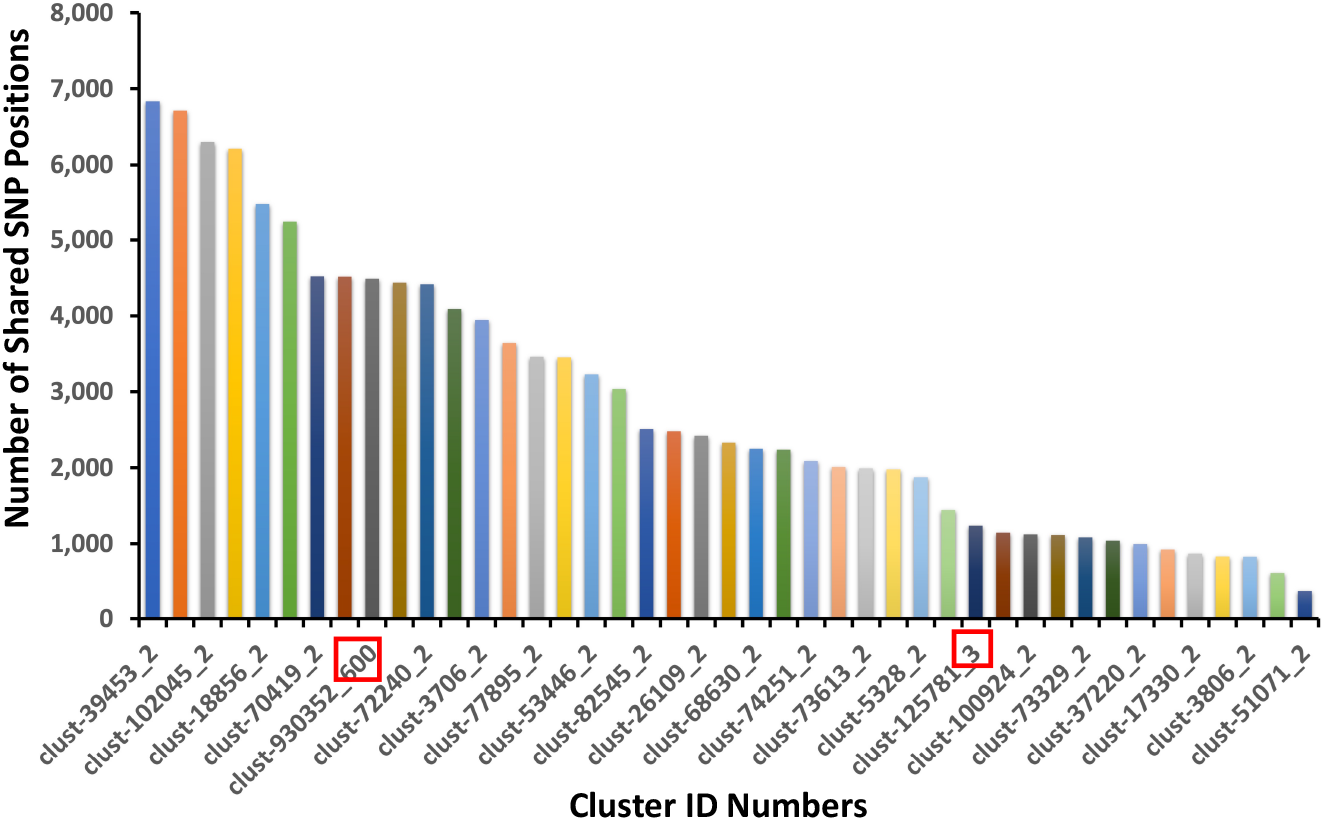
The detection of groups of mutants with shared variants using a clustering algorithm. A total of 930,352 clusters were identified, and these contained an average 78 mutant individuals per a cluster, and an average of 1.3 shared mutation positions per cluster. The top 43 clusters contained an average of 2,831 shared mutation positions, with 41 of the clusters defined by only pairs of mutant individuals. The number of mutant individuals in a cluster is appended to the end of the cluster ID name. Cluster clust-930352_600 (shown in red box) contained 4,487 mutation positions shared by all 600 mutants. Cluster clust-125781_3 (shown in red box) also contained 1,238 mutations shared by three mutants.

### SNP Impact Annotation

Using the snpEff software, we annotated 10,263 distinct high impact SNP positions affecting 7,979 distinct sorghum genes. This includes 7,217 stop-gained (nonsense) mutations and represent averages of 17 high impact SNPs and 12 stop-gained mutations per mutant (Table 3). A total of 136,639 distinct missense SNPs were predicted to have a moderate impact on gene function in 29,835 distinct sorghum genes, and this represents an average of 228 missense mutations per mutant. A total of 83,166 SNPs were synonymous (silent) substitutions. Overall, 30,285 distinct sorghum genes were impacted by moderate-to-high impact SNPs. The remaining SNP effect annotations include 293,332 and 2,884,020 SNPs derived from introns and intergenic regions respectively (Supplemental File S6). Using the SIFT program and the UNIREF90 protein database, analysis of the non-distinct 291,043 missense SNPs (overlapping various isoforms of gene transcripts), showed 106,520 were likely deleterious, 90,892 were likely tolerated, while the remaining 93,631 were indeterminate (Supplemental File S7). The detailed annotation of all the high and medium impact SNPs, including the description of the impacted genes has been included in the analysis (Supplemental File S8).

**Table 3.** Summary predicted impact of likely EMS-induced SNPs on gene function for the 600 sorghum BTX623 mutants

| **SNP Impact Type** | **Total** | **Mean** |
| --- | --- | --- |
| **High Impact** | 10,263 | 17 |
| **Stop Gained** | 7,217 | 12 |
| **Splice Site Donor** | 1,370 | 2 |
| **Splice Site Acceptor** | 1,432 | 2 |
| **Stop Lost** | 19 | 0 |
| **Start Lost** | 226 | 0 |
| **Moderate Impact** | 136,639 | 228 |
| **Missense** | 136,639 | 228 |
| **Low Impact** | 98,623 | 164 |
| **Silent** | 83,166 | 139 |
| **Start Gained** | 7,491 | 12 |
| **Intron variant** | 293,332 | 489 |
| **Intergenic Region** | 2,884,020 | 4,807 |
| **Modifier** | 3,417,516 | 5,696 |

## Discussion

Although several mutant resources are currently available for sorghum genetics (15,16,78), gene function discovery in sorghum and other crop species would greatly benefit from improvements in computational techniques for the precise detection of DNA mutations ^61^. Several techniques had been suggested for filtering false positives, and these include subtraction of shared variants, applying prescribed variant-quality score thresholds, and the removal of variant calls associated with either extreme strand-bias reads or excessive read coverage depth (43-48, 79). The inappropriate application of these best practices could result in the exclusion of bona fide DNA variants as false negatives or the retention of false-positive detections. As an example, in our previous studies, using the variant quality-score threshold of 20 in the initial filtering of SNPs showed marginal improvement (16,80,81), while in another study a quality-score threshold of 100 was effective in the detection bona fide fast neutrons-induced mutations in rice (20). In the specific case of EMS mutagenesis, the likelihood of detecting the same mutation in two or more independently mutagenized individuals in a population is very low, since EMS action impacts random genomic positions. Also, in comparison to other mutagens such as fast neutrons, EMS mutagenesis results in predominantly G/C to A/T DNA transitions (68,69) and hence is limited to a lower diversity in categories of genome target sites (16,19,20). However, the well-known EMS mutation spectrum can be leveraged in determining the suitability or effectiveness of any of the above-mentioned false-positive variant filtering techniques for specific studies. Guided by these well-described characteristics, and the expected mutation spectrum of EMS we proposed and implemented a high-resolution method (Figure 1) for detecting induced mutations in the low-coverage NGS sequencing data for the M_3_ generation from 600 EMS-mutagenized sorghum individuals (Table 1). Using the proposed method, we empirically determined the phred-scale variant quality-score threshold precisely when the retained SNPs in the mutant population were predominantly G/C to A/T substitutions (Figure 2). Using the proposed method we predicted 3,274,606 likely EMS-induced SNPs in the 600 mutant individuals (Table 2 and Figure 3) of which 96% (3,141,908) were G/C to A/T substitutions as expected by EMS action. This result represents an 87% increase (1,521,203 additional SNPs) over the previous analysis which detected 1,753,403 likely EMS-induced mutations (of which 96.7% were G/C to A/T substitutions) in the same mutant population (16). The previous study was based on analysis using the sorghum reference genome version 2.1, while the current analysis used genome version 3.1. To control for genome version being the source for the increased number of mutations predicted, we re-analyzed the entire 600 genomes by using the reference genome version 2.1. The re-analysis for reference genome version 2.1, similarly predicted 3,289,616 likely EMS-induced SNPs including 3,182,253 (96.7%) G/C to A/T substitutions (Table 2 and Figure 3). Coincidentally, the previous study using version 2.1 of the reference genome also predicted 96.7% (1,695,973 out of 1,753,403) were G/C to A/T substitutions (16). Hence the increase in number of SNPs detected is due to the proposed (improved) method and not the improved reference genome assembly version. This high-resolution analysis also suggests that using a phred-scale variant quality-score threshold of 20 for SNPs filtering in the previous study may have been too conservative and probably caused widespread false-negatives. The new findings from the proposed method were primarily based on the SAMtools variant-calling algorithm. To determine the general applicability of the proposed method, we implemented the methodology by using GATK variant-calling algorithm and re-analyzed the NGS sequencing datasets for the whole mutant population. Using GATK with single individual genotyping, 92.4% (3,248,054) of the 3,514,943 SNPs were G/C to A/T mutations (Table 2; Figure 3 and Supplemental Figure S1). Similarly, using GATK with joint genotyping 93.5% (3,211,794) of the 3,435,790 likely EMS-induced SNPs were G/C to A/T mutations (Table 2; Figure 3 and Supplemental Figure S2). A previous benchmark study using high-confident variants showed ∼92% concordance among three variant callers using Illumina exome sequencing datasets (82). Our results from both GATK- and SAMtools-based algorithms also showed a high-concordance, with 3,074,759 likely EMS-induced SNPs consistently detected in all three implementations (Figure 4). Our analysis similarly revealed a high concordance with 94% of SAMtools SNPs overlapping GATK SNPs at a whole-genome mutant population scale. Despite the high similarity in results, there was a striking marked difference in the initial filtering between SAMtools and GATK. The initial subtraction of shared variants had an immediate desirable impact on the GATK-based approaches, with 88% and 90% (3,412,446 and 3,356,788) of retained unique SNPs being G/C to A/T substitutions (Table 2). In comparison, only 65.4% (4,166,924) of the retained, non-shared SAMtools SNPs were G/C to A/T transitions (Table 2). Also, estimating the phred-scale variant quality-score threshold for SAMtools was relatively straightforward, and resulted in a slightly higher percentage of G/C to A/T substitutions (96% versus 94%). We also observed that the vast majority of GATK-specific SNPs which were not predicted by SAMtools, were less likely to be EMS-induced mutations (Supplemental Table S1). These differences in characteristics reflect the underlying assumptions in the two variant-calling algorithms (39,83). Although subtraction of shared variants is a very potent technique, it could inadvertently result in bona fide false-negative mutations being removed. Pollen contamination is quite often a source of shared variants in a sequenced population. Previous attempts to recover false-negative shared variants involves retaining shared G/C to A/T substitutions with high variant-quality score that were detected in up to two mutants (16). We examined the EMS mutation spectrums for various categories of initially predicted SNPs. We observed that among the various SNP categories the G/C to A/T substitution percentage was highest for shared SNPs detected in pairs of mutants; even higher than the percentage for unique SNPs (68 % vs 65%; Supplemental Figure S3). We implemented a clustering algorithm for recovering false-negative shared variants in the mutant population. The algorithm uncovered 42 clusters containing 235,736 likely false-negative SNPs, of which 98.7% were G/C to A/T substitutions. Using the clustering algorithm, we were able to determine that at least 232,022 of 661,156 initially subtracted (shared) SNPs originating from pairs of mutants, and only 3,714 of 291,966 initially subtracted (shared) SNPs detected in triplets of mutant individuals, are most likely bona fide EMS-induced mutations (Figures 5 and 6). Previous studies had shown SNPs associated with extreme strand-biased reads as a potential source of false-positives in Illumina NGS sequencing data (84). Next, we investigated the evidence of extreme strand-bias impact in the filtered SNPs dataset. Of the 307,434 final SNPs having least 95% of the reads derived from a single strand, 94% (290,110 SNPs) were G/C to A/T substitutions, and most likely EMS-induced mutations (Figure 5). This suggests that for low-coverage whole-genome NGS sequencing datasets, the lack of sufficient read depth might result in artificial extreme strand-biased reads mapping at some SNP positions, this might result in these SNPs being discarded as false positives. The snpEff annotation of the final 3,497,654 SNPs predicted 10,263 distinct high-impact and 136,639 moderate impact SNPs in the sequenced mutant population (Table 3). This represents an average of 17 high impact and 228 moderate impact mutations per mutant line. Previous studies revealed that known variants that alter phenotypes usually corroborate their earlier prediction as deleterious mutations, and as such computational predicted deleterious mutations may likely alter phenotypes ^85^. Using the UNIREF90 reference proteome dataset, our analysis predicted 106,520 missense SNPs to be likely deleterious, 90,892 as likely tolerated, while the impact of the remaining 93,631 missense SNPs could not be reliably determined. In summary, we have created an improved large-scale mutant resource for sorghum genetics and created a light-weight web portal for querying the mutant database and seamless link to requesting seed stock for the collections in the Germplasm Resources Information Network (GRIN) database.

## Materials and Methods

### NGS Sequence Retrieval and Reads Mapping

The Illumina next-generation sequencing data for the M_3_ generation from 600 EMS-mutagenized sorghum mutants were retrieved from the NCBI Sequence Read Archive (SRA Accession Number SRP065118) by using the *fastq-dump* program (version 2.9.0) with non-default parameters: “--split-3” (70,71). We retrieved the reference genome sequences and annotations for the sorghum BTx623 genome assembly versions 2.1 and 3.1, from the Phytozome web portal (72). Prior to NGS reads mapping, the reference genome sequences were indexed using *index* command in the BWA short reads aligner (version 0.7.15) (73). The NGS reads for each sequenced sample were mapped to the sorghum reference genome using the BWA *mem* command with non-default parameter: “-t 64”.

### SAMtools-Based Variant-Calling

The SAMtools variant-calling was performed using version 1.8 of the software package (38). Prior to variant-calling, the sorghum reference genome sequences were indexed using the SAMtools *faidx* command with default parameters. The BWA-mapped NGS reads were sorted using the SAMtools *sort* command with non-default parameters: “-@ 64 -m 2G -O bam -T”. Duplicate NGS reads were identified by using GATK *MarkDuplicates* command (version 4.0.5.0) with default parameters (41). The resulting BAM-formatted files were indexed using the *index* command with non-default parameters: “-b -@ 64”. The genotype likelihoods were computed using the BCFtools *mpileup* command (version 1.8) with non-default parameter: “-Ou”. Final variant detection was performed using the BCFtools *call* command with non-default parameters: “-vmO z”. Based on the SAMtools zygosity call encoding in the output variant call files SNPs were characterized as either homozygous or heterozygous.

### GATK-Based Per-Sample Single Individual Variant-Calling

For the GATK-based variant-calling the sorghum reference genome sequences were indexed using the GATK *CreateSequenceDictionary* command (version 4.0.5.0) with default parameters (39). Using the above BWA-sorted and indexed NGS reads, the initial GATK single sample variant-calling for single individuals was performed by invoking the GATK *HaplotypeCaller* with non-default parameters: ‘--java-options “-Xmx4g” HaplotypeCaller’.

### GATK-based Joint Genotyping

For joint genotyping with GATK, the initial steps were similar to the above described GATK approach for per-sample individual variant calling, with the exception of invoking the GATK *HaplotypeCaller* with non-default parameters: ‘--java-options “-Xmx4g” HaplotypeCaller -ERC GVCF’. Next, we created two additional input files: the first was the map file which contained the sequenced sample ID names and their corresponding individual genotyping call filenames. The second file contained the list of chromosome and super scaffold names extracted from the reference genome sequence assembly file. Specifying these two input files, we executed the GATK *GenomicsDBImport* command with non-default parameters: ‘--java-options “-Xmx4g” --batch-size 100 --reader-threads 50’ to merge the individual sample variant call files, which were then subsequently imported into a *GenomicsDB* workspace. Joint genotype calling was performed using the GATK *GenotypeGVCFs* command with non-default parameters: “-G StandardAnnotation -new-qual”.

### Variants Filtering

For both SAMtools-based and GATK-based approaches, all the initially predicted SNPs for the 600 mutants were combined and sorted by their genomic positions. Using this information all variants detected at the same genomic position in multiple mutant individuals were subtracted (44). We calculated the mutation spectrum for the retained unique SNPs to check the effect of the likely false-positive subtraction. We next empirically determined the phred-scaled variant-quality score threshold for further SNPs filtering, by using a high-resolution binning method as follows. All the retained SNPs from the shared variants subtraction were sorted into bin categories corresponding to their respective variant-quality scores, and as such SNPs with the same quality-score were assigned to the same bin category. With emphasis on the percentage of mutations that were G/C to A/T substitutions, we plotted the mutation spectrum for each quality-score bin category. Visual inspection by starting from the lowest to the highest phred-scaled quality-score bin category, we determined the threshold variant quality-score, at which the percentage of SNPs that were G/C to A/T transitions approached the expected results of EMS action. The subset of SNPs with variant quality-score less than the threshold score were filtered. We separately estimated the SNPs quality-score thresholds for the SAMtools- and GATK-based variant-calling results.

### Detection of Likely False-Negative Variants

We implemented a clustering algorithm for detecting and recovering likely false-negative shared variants that were subtracted in the initial SNPs filtering step. We hypothesize that if a set of mutants are genetically related, then a sufficiently large number of (shared) mutations would be detected exclusively in the specific set of mutants and absent in the remaining unrelated mutants. All the initial SAMtools SNP calls in each variant call file were tagged (prefixed) with their respective sample ID names. Next, the sample ID-tagged SNPs from all 600 mutants were combined into a single file. The combined file was sorted by using the genome positions of the SNPs, and the sorted file was used in creating a summary table including the following three column entries: *genome position, number of samples*, and *list of sample IDs*. For example, the table entry “*Chr04:23080886 3 SRR2759335*|*SRR2759504*|*SRR2759646*” means the SNP at genome position 23,080,886 on chromosome 4, was detected in three mutants with sample ID names *SRR2759335, SRR2759504*, and *SRR2759646*. Each grouping of sample IDs on each line in the summary table was treated as a candidate cluster of related mutants. For each candidate cluster, we extend the search by examining the rest of the file to count how many times the same n-tuple of mutants occur exclusively together (i.e., determine the remaining set of mutations that are shared exclusively by the specific set of mutants). For each candidate cluster we recorded the number of shared mutations and the percentage of mutations that were G/C to A/T substitutions. Next, the detected clusters were sorted and ranked based on their number of shared mutations. The top-ranked clusters with markedly high number of shared mutation counts were retained and their corresponding percentages of G/C to A/T substitution was used as a benchmark evidence of likely EMS-induced origins. The identified, likely false negatives were then added to the unique (non-shared) filtered SNPs. For each affected mutant individual, the merging of the likely false-negatives and the likely true-positives was accomplished by appending the shared variants file to their corresponding filtered unique variants file and re-sorting the resulting variant call file by using the BEDTools *sort* command (version v2.26.0) with the non-default parameter “-faidx chromNames” (74).

### SNP Impact Annotation

We used the snpEff variant annotation package version 4.3t (75) to predict the impact of the detected SNPs on gene function. We created a custom snpEff database for the sorghum reference genome version 3.1 as follows. We edited the snpEff configuration file to include an entry for sorghum genome version 3.1. Next, we added the sorghum genome reference sequences, genome annotation .gff files and the protein sequences to the required subfolders in the snpEff package, and we run the snpEff *build* command with non-default option “-gff3 -v Sbicolor_454_v3.0.1”. Using the variant call files as input, the variant impact annotation was performed by using the snpEff *eff* command.

### SIFT Annotation of Missense SNP Effect

For the final set of retained SNPs we selected all missense substitutions, including the genome position and the DNA base substitution information from their corresponding variant call files. Next, we retrieved the subset of sorghum protein sequences which contained the predicted missense mutations. We next downloaded the UNIREF90 set of protein sequences (109,653,977 sequences in release version 2020_02; 22-Apr-2020) from the UniProt web portal (76). Using the impacted sorghum protein sequences and the UNIREF90 FASTA sequences as query and database options respectively, we used the SIFT4G algorithm (77) with non-default parameter “-t 64”, to predict the likelihood of the missense substitutions having a tolerated or deleterious effect in the genome. Based on the SIFT analysis results, each missense variant was assigned one of the following designations: *TOLERATED, DELETERIOUS* or *UNKNOWN*. For all SNPs with predicted moderate or high impact on gene function, we appended the SIFT predictions, and the gene function description from the sorghum reference genome version 3.1.

## List of abbreviations

SNPs: Single Nucleotide Polymorphisms
GRIN: Germplasm Resources Information Network
NGS: Next-Generation Sequencing
EMS: Ethyl Methanesulfonate
ENU: N-ethyl-N-nitrosourea
MNU: N-methyl-N-nitrosourea
SRA: Short Reads Archive

## Declarations

### Ethics approval and consent to participate

Not Applicable

### Consent for publication

Not Applicable

### Availability of data and materials

All data generated or analyzed during this study are included in this published article (and its supplementary information files).

The NGS datasets for the sequenced sorghum mutants are available in the NCBI SRA repository (Study Accession Number: SRP065118; BioProject ID: PRJNA297450).

Supplemental data files are available via the DRYAD repository (https://doi.org/10.5061/dryad.hmgqnk9hj).

The described sorghum mutations can be accessed at the following link: (http://heliamphora.lcsc.edu/sorghumgenetics2.0/)

## Authors’ contributions

C.A.-Q, C.W, M.T and E.M.B designed the experiments. J.M.S, M.Y.B, A.T.S, D.A and C.A.-Q, performed the variant-calling and annotation. T.C.H implemented the webserver and Y.K implemented the databases. C.K, D.L and C.A.-Q performed the clustering analysis. C.A.-Q drafted the manuscript. C.A.-Q, E.M.B, C.W, and M.T edited the manuscript.

## Acknowledgements

The authors would like to thank the following Lewis-Clark State College administrators for their research support: Jane Finan, Matthew Johnston, Heather Henson-Ramsey, Jerry Hindberg, Lori Stinson, and Martin Gibbs.

## Supplemental Information

### Supplemental Figures

**Supplemental Figure S1.**
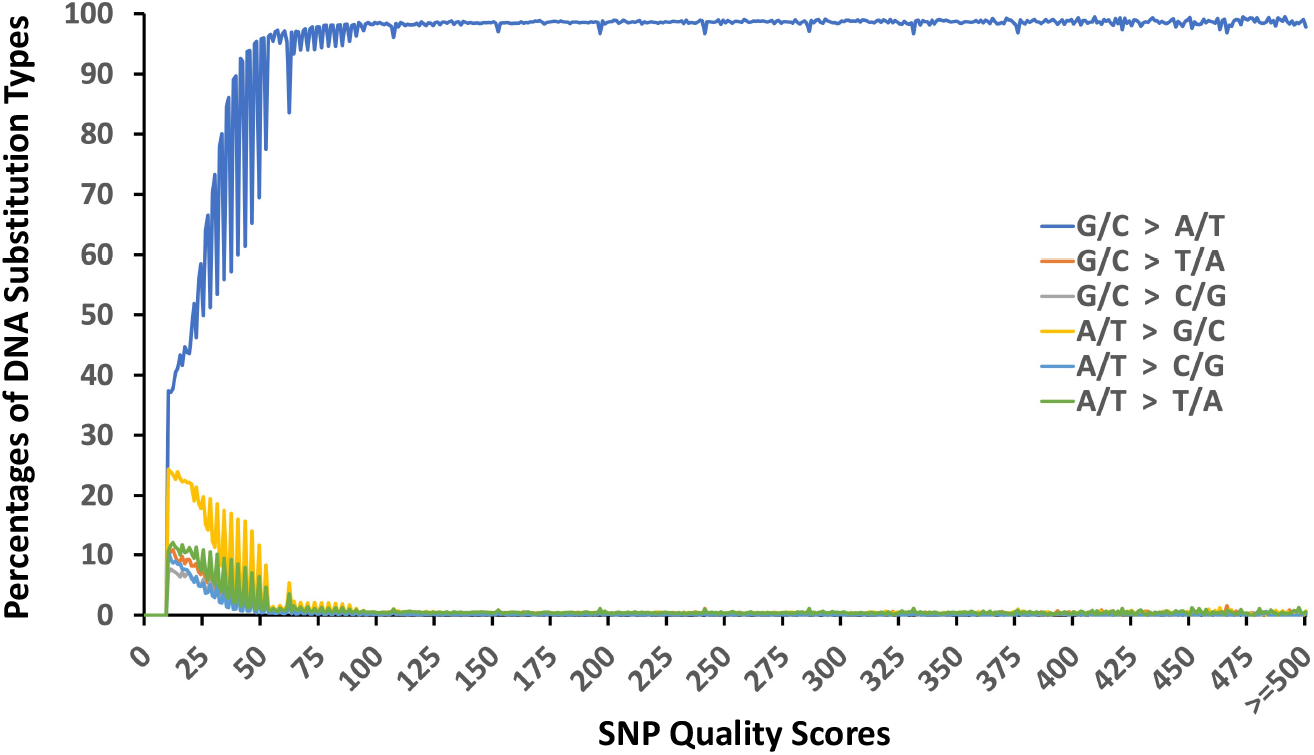
The binning approach to computing the variant quality-score filtering threshold for the initial GATK-predicted single individual genotyping mutations in the 600 sorghum BTx623 mutants. The initially predicted SNPs are assigned to bin categories corresponding to their variant-quality score. The EMS mutation spectrum for all SNPs in each bin category is calculated. At a variant-quality threshold of 26, the percentages of non-G/C to A/T DNA substitutions decrease drastically, while G/C to A/T mutations become the dominant mutation type.

**Supplemental Figure S2.**
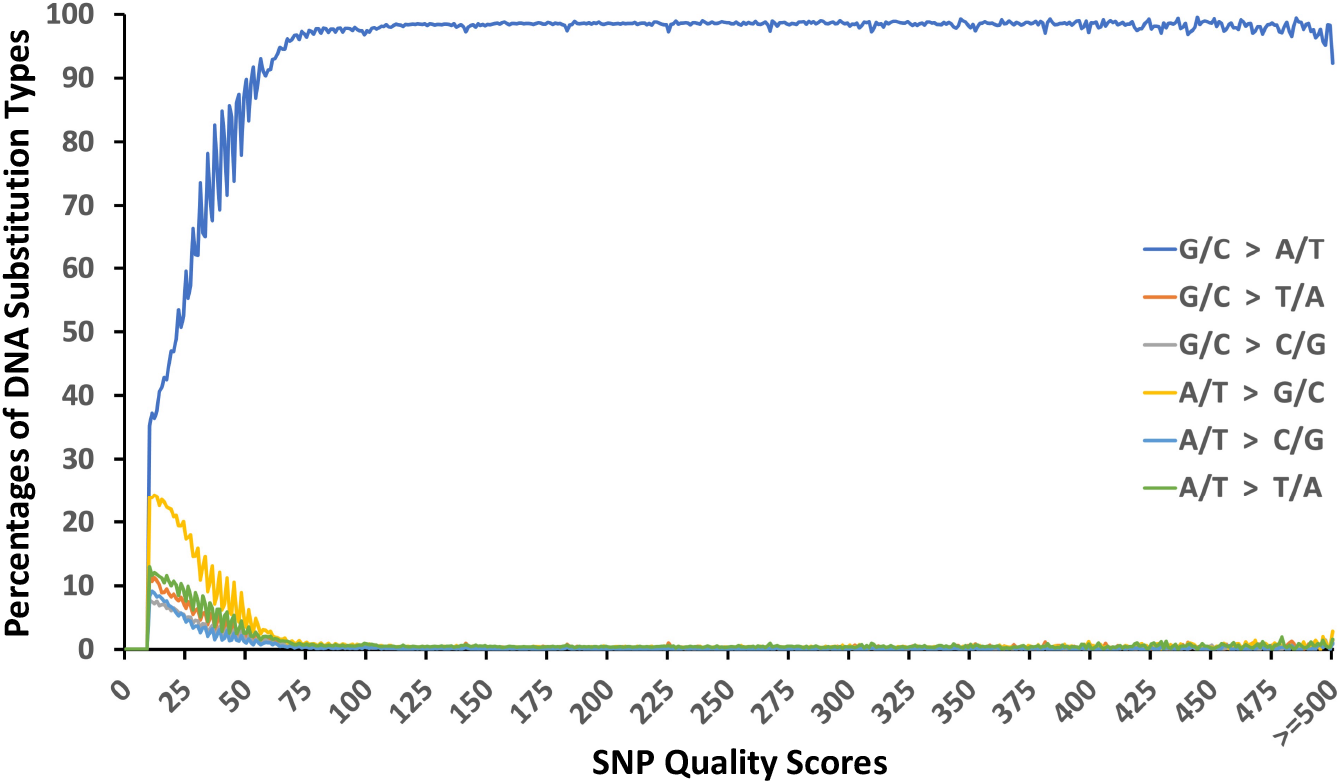
The binning approach to computing the variant quality-score filtering threshold for the initial GATK-predicted joint genotyping mutations in the 600 sorghum BTx623 mutants. The initially predicted SNPs are assigned to bin categories corresponding to their variant-quality score. The EMS mutation spectrum for all SNPs in each bin category is calculated. At a variant-quality threshold of 28, the percentages of non-G/C to A/T DNA substitutions decrease drastically, while G/C to A/T mutations become the dominant mutation type.

**Supplemental Figure S3.**
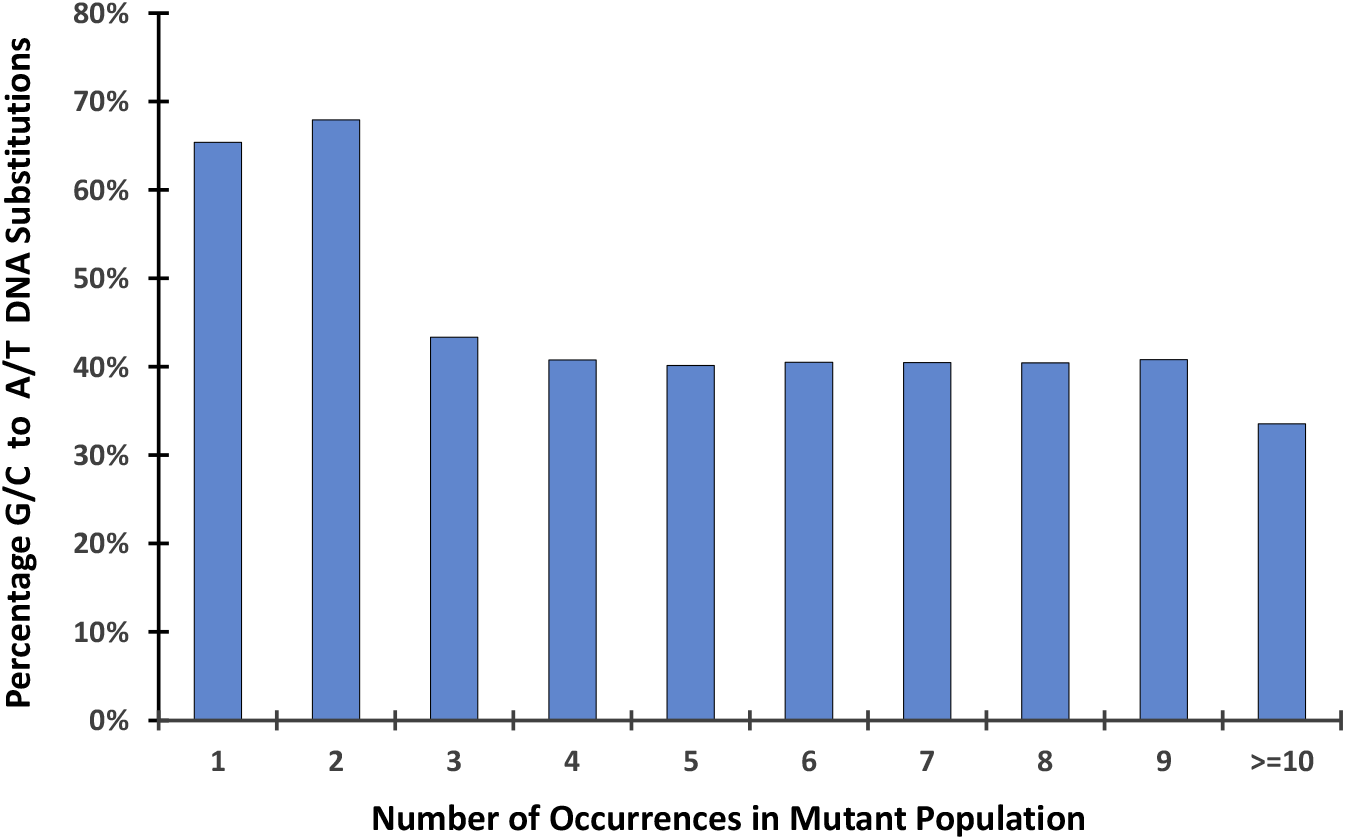
Categories of initially predicted SNPs and their corresponding mutation spectrums. The mutation categories are based on the number of times a mutation is detected in the mutant population. For example, category 1 and category 2 refer to mutations detected once and twice, respectively in the mutant population. The percentage of G/C to A/T DNA substitutions for category 2 is unexpectedly higher than category 1.

### Supplemental Tables

**Supplemental Table S1.** The mutation spectrums for the final filtered GATK-specific SNPs that were not detected by SAMtools.

|  | GATK-Single<sup>a</sup> | GATK-Joint<sup>b</sup> |
| --- | --- | --- |
| G/C > A/T | 232,234 | 207,491 |
| G/C > T/A | 32,465 | 26,505 |
| G/C > C/G | 22,990 | 18,618 |
| A/T > G/C | 74,779 | 58,456 |
| A/T > C/G | 21,043 | 16,461 |
| A/T > T/A | 39,376 | 32,380 |

<sup>a</sup>GATK SNPs-calling with single individual genotyping.
<sup>b</sup>GATK SNPs-calling with joint genotyping.

